# Regional and Global Analysis of Cerebrovascular Reactivity Using 4D Flow MRI

**DOI:** 10.1101/2025.06.23.661174

**Authors:** Sergio Dempsey, Harvey Walsh, Debbie Zhao, Gina Quill, Martyn P. Nash, Kelly Burrowes, James P. Fisher, Gonzalo D. Maso Talou

## Abstract

Cerebrovascular reactivity (CVR) assessments are promising for diagnostic and prognostic applications related to vascular function and neurodegenerative diseases. Current vascular imaging-based CVR indices have been limited to single-vessel analysis and report conflicting results, for example, whether there are differences in CVR between sexes. Alternative vascular imaging-based indices can be derived from 4D flow MRI, namely intracranial pulse wave velocity and pulsatility transmission, which are based on vascular regions and offer a more comprehensive perspective on vascular function that can also scale to a global level. To date, these types of indices have not been measured with a vascular stimulus. In this work, standard single-vessel and novel regional and global CVR indices were studied in 10 healthy young adults (29 *±* 2 years, 5 male, 5 female) in response to hypercapnia. Consistent increases in flow, area and velocity manifested at all levels and were greater in males. Regional pulse wave velocity and pulsatility transmission offered a minimum 40% increase in the dynamic range of CVR magnitudes that may benefit cohort stratification. Notably, pulse wave velocity increased in 50% of the vascular regions (most probably due to vasoconstriction), accompanying an increase in flow; a profound observation given the common expectation of vasorelaxation with hypercapnia. Consistently reduced flow, velocity, area, pulse wave velocity, and transmission index reactivities in females may partially explain the varying pathological outcomes between sexes, evidencing the continued value of 4D flow for the evaluation of CVR in other cohorts.

**New and Noteworthy:** Cerebrovascular reactivity (CVR) of 4D flow-based intracranial pulse wave velocity and pulsatility transmission were evaluated during hypercapnia for the first time. Remarkably, pulse wave velocity increased in some cases and decreased in others, suggesting that vasorelaxation may not always occur during hypercapnia. These indices also expanded CVR dynamic range by more than 40 % compared to common velocimetry techniques, which could offer improved cohort stratification. Evidence of diminished CVR in females was also observed.

## 1 Introduction

Assessment of arterial haemodynamics has become a promising tool for the prognosis of several cerebral pathologies such as small vessel disease and dementia [1], where, in many cases, an elevated arterial pulsatility is a hypothesised initiator of pulse wave encephalopathy [2], [3]. Increased pulsatility can originate from a decreased ability to dampen the incoming pressure wave [4], which is partially regulated by cerebrovascular tone (CVT). Thus, the assessment of CVT is possibly beneficial but currently difficult to characterise. Commonly measured single-site assessments of flows, velocities, or regional perfusions do not fully characterize how CVT actively modulates the pressure wave across the brain. Instead, the change in these hemodynamics must be comprehensively assessed at multiple sites to begin to assess the role of CVT. 4D flow MRI captures the haemodynamics of larger cerebrovasculature, offering several more comprehensive indices that may better characterise CVT and vascular function at the whole-organ level. Pulsatility transmission (*p*_*t*_) [5] and damping (*p*_*d*_) [4] track the change in pulsatility-index (*p*_*i*_) amplitude, identifying how the vasculature and CVT modulate the pressure wave. Additionally, it is possible to measure pulse wave velocity (PWV) [6]–[8], which theoretically reflects effective vascular stiffness, a consequence of CVT. While these regional indices offer a new perspective on vascular function, they still do not demonstrate CVT modulation, which in response to vasoactive stimuli is termed cerebrovascular reactivity (CVR). To our knowledge, the reactivity of region-based PWV and *p*_*t*_ indices have not been demonstrated.

A common and well-studied stimulus to evaluate CVR is hypercapnia [9]. Elevating the inspired fraction of carbon dioxide (CO_2_) raises the concentration of arterial CO_2_ [9] and increases cerebral blood flow (CBF) by eliciting several mechanisms of CVT relaxation [10], [11]. However, hypercapnia-induced CVR is subject to many systemic confounders. Circulating sex hormones play a role in resting CVT and blood flow [11], and may affect CVR, although studies are contradictory (see selected studies with hormone-CVR effects [12]–[14], and others without [15], [16]). Through cardiac and perivascular innervation, chemosensitivity of the carotid body, heart, and peripheral vasculature [17] increases cardiac output [18], systolic blood pressure (SBP) [19]–[21], and diastolic blood pressure (DBP) (indicating peripheral vasoconstriction) [22]. These changes suggest altered cerebral inlet pressure and peripheral resistances, which can impact CVR measurements. Cardiovascular chemosensitivity has also been shown to be sensitive to sex hormones, with testosterone receptors present in the carotid body [23], and a greater blood pressure response during the luteal versus follicular phases of the menstrual cycle [24]. Additional confounders include age and health status [9], and possibly sex, with some studies showing females are more reactive than males, while others demonstrate the opposite (see [25] and references therein).

To demonstrate the feasibility and preliminary CVR of whole-organ measurements characterised by 4D flow MRI region-based *p*_*t*_, *p*_*d*_, and PWV, a hypercapnia study was designed to control for the effects of age and pathology with near simultaneous measurements of sex hormones, blood composition, and blood pressure. To assess region-based CVR indices, their magnitudes were analysed and compared with their vessel-specific and global index counterparts across subjects and vascular regions. Finally, a preliminary covariate analysis of blood and cardiac factors with CVR was explored to assess the strength of confounder influence.

## 2 Methodology

### 2.1 Ethics Statement and Participant Description

The “Haemodynamic Encephalopathy Risk (HER) study” is a prospective study to explore sex and ethnicity differences in CVR in young healthy individuals in New Zealand. The study was approved by the Central New Zealand Health and Disability Ethics Committee (HDEC-20417), registered under the Australian New Zealand Clinical Trials Registry (ACTRN12624001097538), and performed in accordance with the guidelines of the 1964 Declaration of Helsinki and its later amendments. All participants provided written informed consent.

Ten participants (5 male, 5 female) were recruited (aged 29 *±* 2 years, range [26 ™ 34] years). All participants were non-smokers, normotensive, and of European descent. Exclusion criteria were: history or evidence of vascular disease, functional or structural cardiac abnormalities (e.g., atrial fibrillation or septal defect), diabetes, contraindications to MRI or blood sampling, brain pathology (e.g., cerebrovascular disease or chronic headaches), and diagnosed mental health conditions. All MRI scans were reviewed by a neuroradiologist for incidental findings and excluded if structural abnormalities were present.

### 2.2 Experimental Protocol

Participants were scheduled for two visits. Visit 1 was for familiarisation, where demographics were collected and the participants experienced the hypercapnic gas mix (described in the Gas Protocol section) during free inspiration for 15 minutes. Visit 2 was for data collection. Participants abstained from alcohol and exercise 24 hours prior, and fasted for 2 hours prior to arrival. Cerebral MRI, brachial blood pressure measurements and venous blood samples were collected within a 1.5 hour window. The order was dependent on scheduling availability. Approximate protocol timing is provided in Figure 1A as an aid to the following methodological descriptions.

**Figure 1.**
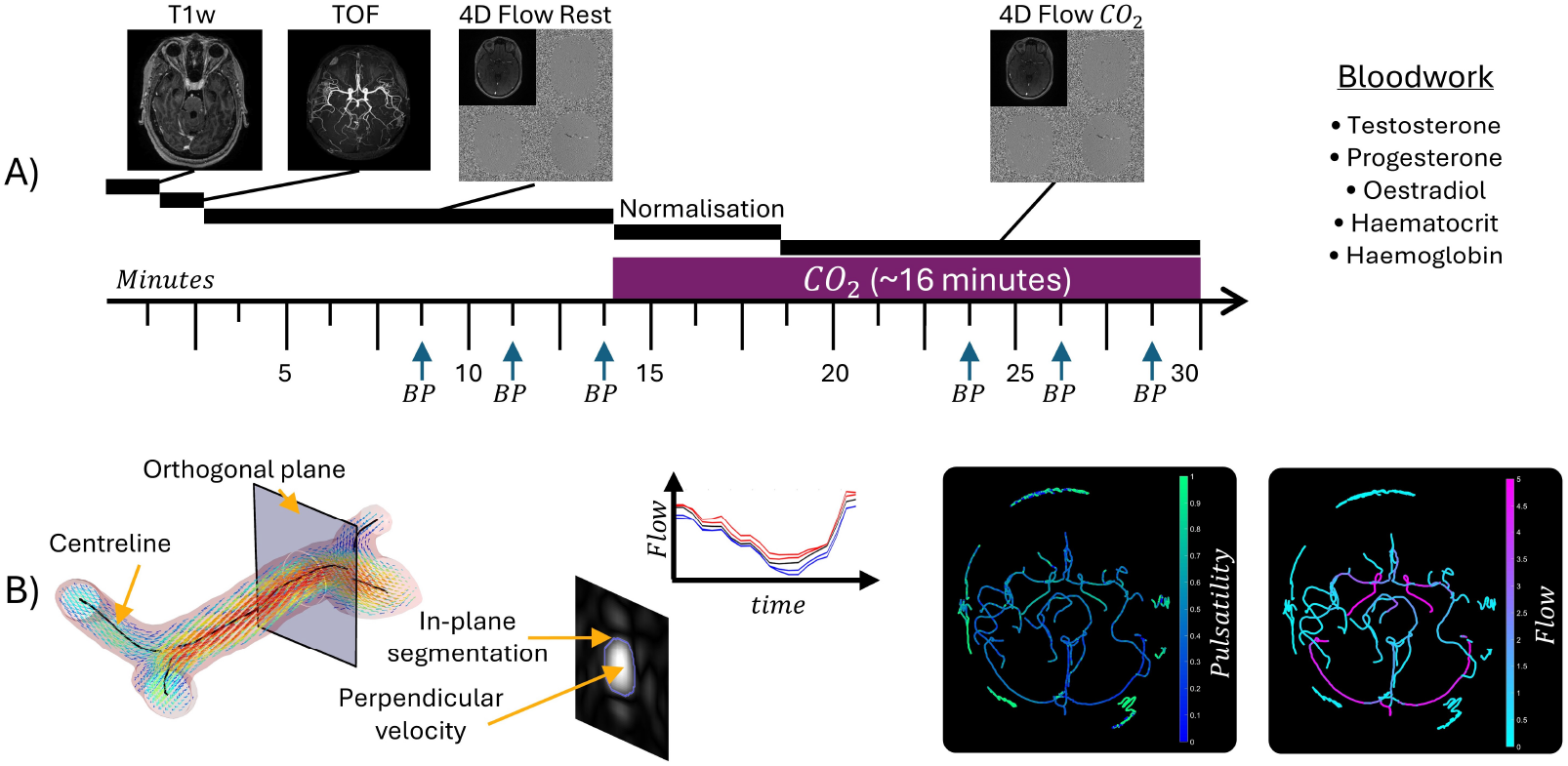
A) Protocol for the Haemodynamic Encephalopathy Risk study demonstrating the order of imaging sequences and estimated timings of scans and blood pressure measurements. B) 4D flow processing workflow to segment vessels, interpolate blood flow at vascular cross-sectional planes, and compute through-plane flow during the cardiac cycle. To the right are visualisations of the spatially varying haemodynamic data throughout the cerebrovasculature. Symbols and abbreviations: blood pressure (BP), carbon dioxide (*CO*_2_), T1-weighted (T1w), time of flight (TOF).

#### 2.2.1 Gas Protocol

CO_2_ enriched gas, mixed on site (GSM-3 Gas Mixer, CWE Inc.), was composed of 4% CO_2_, 21% oxygen and balanced nitrogen. The ratios were confirmed within a tolerance of 0.1% using a CO_2_ and O_2_ gas analyser (ML206, ADInstruments). Gas was administered through a sealed face mask (7450 Series Silicone V2™Oro-Nasal Mask, Hans Rudolph) connected to a two-way, non-rebreathing valve (T-2700, Hans Rudolf Inc.), which was in turn attached to 150 L non-diffusing Douglas bags (ADInstruments). Each bag provided approximately 7 ™ 12 minutes of breathing time depending on the breathing rate and tidal volume of the participant. Several bags were connected in tandem with 0.9 m of tubing between each bag (Hans Rudolph Inc.) to ensure safe supply during the experimental sessions. Gas release was controlled by a manual directional control valve (three-way T-shape stopcock-type™, Hans Rudolph). Partial pressures of CO_2_ were monitored using the gas analyser sampled at 1 kHz and stored using Labchart software (Labchart Pro v8.1.30, ADInstruments).

#### 2.2.2 MRI Protocol

Participants were imaged at the University of Auckland Centre for Advanced MRI (CAMRI) with a 3T scanner (MAGNETOM Vida Fit, Siemens Healthineers) using a 32-channel head coil. A structural T1 magnetisation prepared rapid acquisition gradient echo (MPRAGE) sequence was acquired with the following typical imaging parameters: field of view (FOV) 256×256×256 mm, resolution 1.3 mm isotropic, repetition time (TR) 19.4 ms, echo time (TE) 3.69 ms, flip angle (FA) 20°, phase-encoding generalised autocalibrating partially parallel acquisition factor (peGRAPPA) 4, phase partial Fourier 6*/*8. Then, a coarse 3D gradient echo fast low-angle shot time-of-flight with compressed sensing (CS) angiogram was acquired for 4D flow planning centred on the cranial base: FOV 200×200×156 mm, resolution 0.5 mm isotropic, TR 1590 ms, TE 2.64 ms, TI 880 ms, FA 8°, CS total factor 10.3, Cartesian k-space trajectory. This was followed by a 3D gradient echo 4D flow sequence: FOV 170×224×40 mm (right-left, anterior-posterior, superior-inferior, respectively), resolution 1.0 mm isotropic, TR 72.8 ms, TE 3.34 ms, FA 15°, peGRAPPA 2, phase and slice partial Fourier 6*/*8, Cartesian k-space trajectory, velocity encoding 90 m/s, k-space segments 3, cardiac phases 20 (temporal resolution 48 ± 9 ms) using retrospective cardiac binning estimated from a finger pulse oximeter. After the first 4D flow sequence, a researcher in the scanner room started the gas challenge by switching on the CO_2_ enriched gas. Participants were given *≈* 5 minutes for homeostatic normalisation, after which a repeat 4D flow scan was acquired while the gas remained on. The entire imaging protocol took *≈* 30 minutes. The timings for the start and end of each sequence were tagged in the LabChart recording to identify the gas dynamics during each sequence for post-processing analysis described in the gas analysis section. The gas analyser was in the control room with a 9.15 m sample line connected to the participant’s mask in the scanner room.

#### 2.2.3 Blood Pressure Protocol

Blood pressure of the participants was measured before or after the MRI scan. While resting in a bed in the lateral (left) decubitus position, masked participants were fitted with a brachial pressure monitor (BP+ Supra-systolic oscillometric central blood pressure monitor, Uscom) and rested for a gas exposure with similar timings to the MRI protocol (15 minutes normocapnia, 15 minutes hypercapnia). The gas analyser was beside the participant bed, with a 3.05 m sample line to connect the analyser. During this period, starting at 7 minutes into normocapnia, blood pressure was measured three times with 1 minute between measurements, and the SBP and DBP were recorded. This was repeated again 7 minutes into hypercapnia. The groups of three measurements were averaged to provide SBP and DBP values during normocapnia and hypercapnia.

#### 2.2.4 Blood Test Protocol

Blood was sampled and subsequently analysed at LabPLUS (https://www.labplus.co.nz, less than 5 minutes walk from CAMRI), annually accredited for “good laboratory practice” by International Accreditation New Zealand, ensuring that quality was consistent with international standards (ISO15189:2012, “Medical Laboratories – Requirements for quality and competence”). Analysis consisted of measuring oestradiol, progesterone and testosterone concentrations, and a full blood count, with the measurements of interest being haematocrit (Hct) and haemoglobin (Hgb). Hgb was evaluated due to its role in gas dynamics and transport.

### 2.3 Data Analysis

#### 2.3.1 Gas Analysis

Using a Labchart waveform analysis module, the end-tidal partial pressure of carbon dioxide (P_*ET*_ CO_2_) were automatically identified for each respiratory cycle. The P_*ET*_ CO_2_ time series was then exported for further processing in Matlab (v2024a, Mathworks), where each of the time series were filtered for outlier removal using a Hampel filter with a window length of 20 seconds and an outlier exclusion of 1.5*σ* [26]. Then, using the sequence time tags from the Labchart recording, the mean P_*ET*_ CO_2_ values during normo- and hypercapnia were computed over the 4D flow acquisition times.

#### 2.3.2 MRI Analysis

The 4D flow data were processed using the Quantitative Velocity Tool (QVT) pipeline in Matlab [27], [28] performing automatic vessel segmentation, background correction and cross-sectional mean flow, area (CSA), mean velocity (CSV), and pulsatility index (*p*_*i*_) analyses (see Figure 1B for a visualisation of QVT processing and spatially computed values of interest). During the initial development of the 4D flow protocol, it was observed that the built-in QVT segmentation algorithm did not provide acceptable segmentation quality for subsequent analysis, so an external vascular segmentation software [29] was used and then the obtained segmentation was loaded to continue the remaining QVT pipeline tasks.

After baseline QVT processing, the QVT+ module was then used to extract all vessel-specific and region-based indices. For vessel-specific analysis, the flow, pulsatility, vessel-specific CSA and CSV of the main circle of Willis arteries were extracted matching previous works [30]–[32]. For regional analysis, the regions considered were the vascular territories originating from the left internal carotid artery (LICA), right internal carotid artery (RICA) and basilar artery (BA). The regional indices derived where *p*_*d*_, *p*_*t*_, PWV, and average CSV and CSA (see Results for visualisation of sampling points and regional territories). All indices were computed as previously described [5], [8]. For global analysis, the region-based indices were averaged to provide global values. In addition, a global CBF was calculated as the sum of the ICA and BA flows, representing the total CBF. For each participant, the total CBF was normalised with reference to their total volume of white and grey matter, which was segmented from T1-weighted sequences using the computational anatomy toolkit (CAT12, v12.8, [33]) in Matlab. In the event that a regional index could not be computed (for instance, PWV can be inconclusive), or a reference regional artery (ICA or BA) was not identifiable for *p*_*t*_ or *p*_*d*_, the remaining regions were averaged. Additionally, the time-averaged heart rate (HR) was extracted from each 4D flow scan, representing an average HR over the acquisition time during normocapnia and hypercapnia.

### 2.4 Reactivity Analysis

#### 2.4.1 Reactivity Estimation

For each index *I*, its reactivity *I*_*r*_ was calculated as

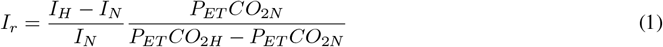

where subscripts *H* and *N* refer to the hypercapnia and normocapnia measurements. During reporting, *I* is replaced by the name of the index, and it is expressed in percent differences (%Δ*/*%Δ).

#### 2.4.2 Assessing Index Reactivity

The sex-specific reactivity of each index was assessed with a one-way Wilcoxon signed-rank test. Sex differences for each index were then evaluated using two-way Wilcoxon rank-sum tests. Index data from different locations were combined for a repeated assessment under the assumption of measurement independence. All statistical analyses were identified as significant for *p <* 0.05.

The number of “confident” CVR responses was evaluated by summing the total percentage of all measurements that were beyond one standard deviation of the mean test-retest variability as reported in the literature or estimated in the present study (see Supplementary Note 1). Given the recent development of regional 4D flow analysis, test-retest variability was not available for all indices. For vessel-specific test-retest variability, the following standard deviations were used: flow *σ* = 16.0% [34], [35], CSA *σ* = 11.5% (see Supplementary Note 1), CSV *σ* = 8.5% [34], [36], and *p*_*i*_ *σ* = 14.5% [35], [37]. For region-based test-retest variability, the following standard deviations were used: PWV *σ* = 8.5% [37], CSA *σ* = 16.5% (see Supplementary Note 1) CSV *σ* = 15% (see Supplementary Note 1), *p*_*d*_ *σ* = 6.6% (see Supplementary Note 1), *p*_*t*_ not available. For global test-retest variability, the average % standard deviation from this cohort during normocapnia was used: P_*ET*_ CO_2_ *σ* = 1.7%, SBP *σ* = 3.1%, DBP *σ* = 5.9%, and HR *σ* = 8.9% (standard deviation of all R-R intervals during the normocapnic 4D flow scan). The total CBF test-retest error was *σ* = 3.7% [37]. Global measures of additional cerebrovascular indices were not available.

#### 2.4.3 Covariate Analysis

Global index correlation strengths were evaluated with other global systemic indices (circulating blood composition data, exhaled gas concentrations, brachial blood pressure) for covariate analysis during normocapnia and hypercapnia.

## 3 Results

### 3.1 Study Execution

All participants scheduled for visit 2 completed the protocol. Concerning the order of imaging and blood work, 7*/*10 participants were scheduled to undergo MRI, blood pressure, and then blood work. For one participant, blood pressure was recorded before MRI, and two participants had blood samples taken before the MRI and blood pressure recordings. The mask of one male participant leaked during the MRI session (identified by the lack of increased CO_2_ during exhalation), resulting in exclusion from the MRI reactivity analysis. Data acquired during normocapnia and the separate blood pressure session for that participant were still included in the analysis.

### 3.2 Baseline Characteristics

As summarised in Table 1, baseline measurements collected for covariate analyses were within nominal ranges, with additional baseline reporting for select cerebrovascular indices. Vessel-specific baseline values are not reported here as they are not used in covariate analysis; for normative values, see [28]. Several statistically significant differences between males and females were observed: females had higher volume-normalised total CBF and lower P_*ET*_ CO_2_, testosterone, Hct, and Hgb. While progesterone and oestradial were expected to be higher in females on average, the large variability of these hormone levels in females meant that statistical significance was not reached.

**Table 1:**
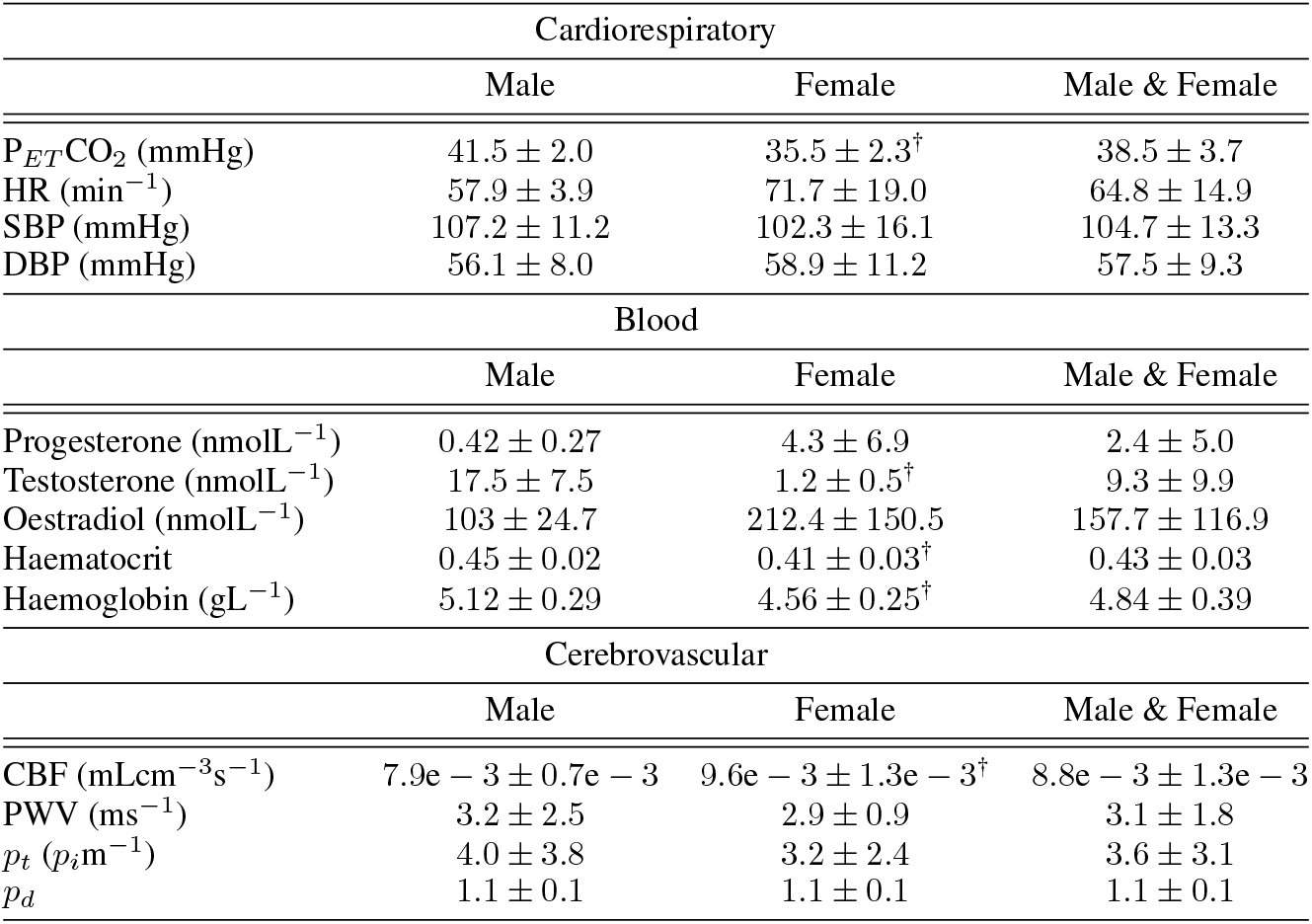
Baseline characteristics for the study cohort. † indicates a statistically significant (*p <* 0.05) sex difference. Symbols and abbreviations: partial pressure of end-tidal carbon dioxide (P_*ET*_ CO_2_), heart rate (HR), systolic blood pressure (SBP), diastolic blood pressure (DBP), cerebral blood flow (CBF), pulse wave velocity (PWV), pulsatility transmission (*p*_*t*_), pulsatility damping (*p*_*d*_).

| Cardiorespiratory |  |  |  |
| --- | --- | --- | --- |
|  | Male | Female | Male & Female |
| $P_{ET}\text{CO}_2$ (mmHg) | $41.5 \pm 2.0$ | $35.5 \pm 2.3^\dagger$ | $38.5 \pm 3.7$ |
| HR ($\text{min}^{-1}$) | $57.9 \pm 3.9$ | $71.7 \pm 19.0$ | $64.8 \pm 14.9$ |
| SBP (mmHg) | $107.2 \pm 11.2$ | $102.3 \pm 16.1$ | $104.7 \pm 13.3$ |
| DBP (mmHg) | $56.1 \pm 8.0$ | $58.9 \pm 11.2$ | $57.5 \pm 9.3$ |
| Blood |  |  |  |
|  | Male | Female | Male & Female |
| Progesterone ($\text{nmolL}^{-1}$) | $0.42 \pm 0.27$ | $4.3 \pm 6.9$ | $2.4 \pm 5.0$ |
| Testosterone ($\text{nmolL}^{-1}$) | $17.5 \pm 7.5$ | $1.2 \pm 0.5^\dagger$ | $9.3 \pm 9.9$ |
| Oestradiol ($\text{nmolL}^{-1}$) | $103 \pm 24.7$ | $212.4 \pm 150.5$ | $157.7 \pm 116.9$ |
| Haematocrit | $0.45 \pm 0.02$ | $0.41 \pm 0.03^\dagger$ | $0.43 \pm 0.03$ |
| Haemoglobin ($\text{gL}^{-1}$) | $5.12 \pm 0.29$ | $4.56 \pm 0.25^\dagger$ | $4.84 \pm 0.39$ |
| Cerebrovascular |  |  |  |
|  | Male | Female | Male & Female |
| CBF ($\text{mLcm}^{-3}\text{s}^{-1}$) | $7.9\text{e} - 3 \pm 0.7\text{e} - 3$ | $9.6\text{e} - 3 \pm 1.3\text{e} - 3^\dagger$ | $8.8\text{e} - 3 \pm 1.3\text{e} - 3$ |
| PWV ($\text{ms}^{-1}$) | $3.2 \pm 2.5$ | $2.9 \pm 0.9$ | $3.1 \pm 1.8$ |
| $p_t$ ($p_i\text{m}^{-1}$) | $4.0 \pm 3.8$ | $3.2 \pm 2.4$ | $3.6 \pm 3.1$ |
| $p_d$ | $1.1 \pm 0.1$ | $1.1 \pm 0.1$ | $1.1 \pm 0.1$ |

### 3.3 Response to Hypercapnia

#### 3.3.1 Vessel-Specific Reactivity

A positive Flow_*r*_ reactivity was identified in almost all vessels (see Figure 2). The Flow_*r*_ reactivity was statistically significant for 1*/*9 male vessels and 4*/*9 female vessels when the data were separated by sex. Combining all vessel data, significant male and female Flow_*r*_ were identified, with significantly greater male Flow_*r*_. Overall, 67*/*73 vessels showed an increase in flow during hypercapnia. The CSA reactivity was less consistent per vessel. Individual vessels did not dilate nor constrict significantly when separated by sex. When combined, both male and female CSA_*r*_ were significantly positive with no sex difference. Overall, 48*/*73 vessels dilated during hypercapnia. Vessel-specific CSV_*r*_ was the most reactive per vessel. 5*/*9 male and 5*/*9 female vessels measured a significant CSV_*r*_, without a difference between the sexes. When combined, both male and female CSV_*r*_ were significant, with significantly higher male CSV_*r*_. Overall, 70*/*73 vessels showed an increase in velocity during hypercapnia. Finally, *p*_*i*_ decreased in response to hypercapnia, with no difference in magnitude between sexes for any vessel. When separated by sex, the decreases in *p*_*i*_ were significant in 2*/*9 male and 2*/*9 female vessels. Assessing combined data, both male and female *p*_*ir*_ were significant with no sex differences. Overall 62*/*73 vessels showed a decrease in *p*_*i*_ during hypercapnia.

**Figure 2.**
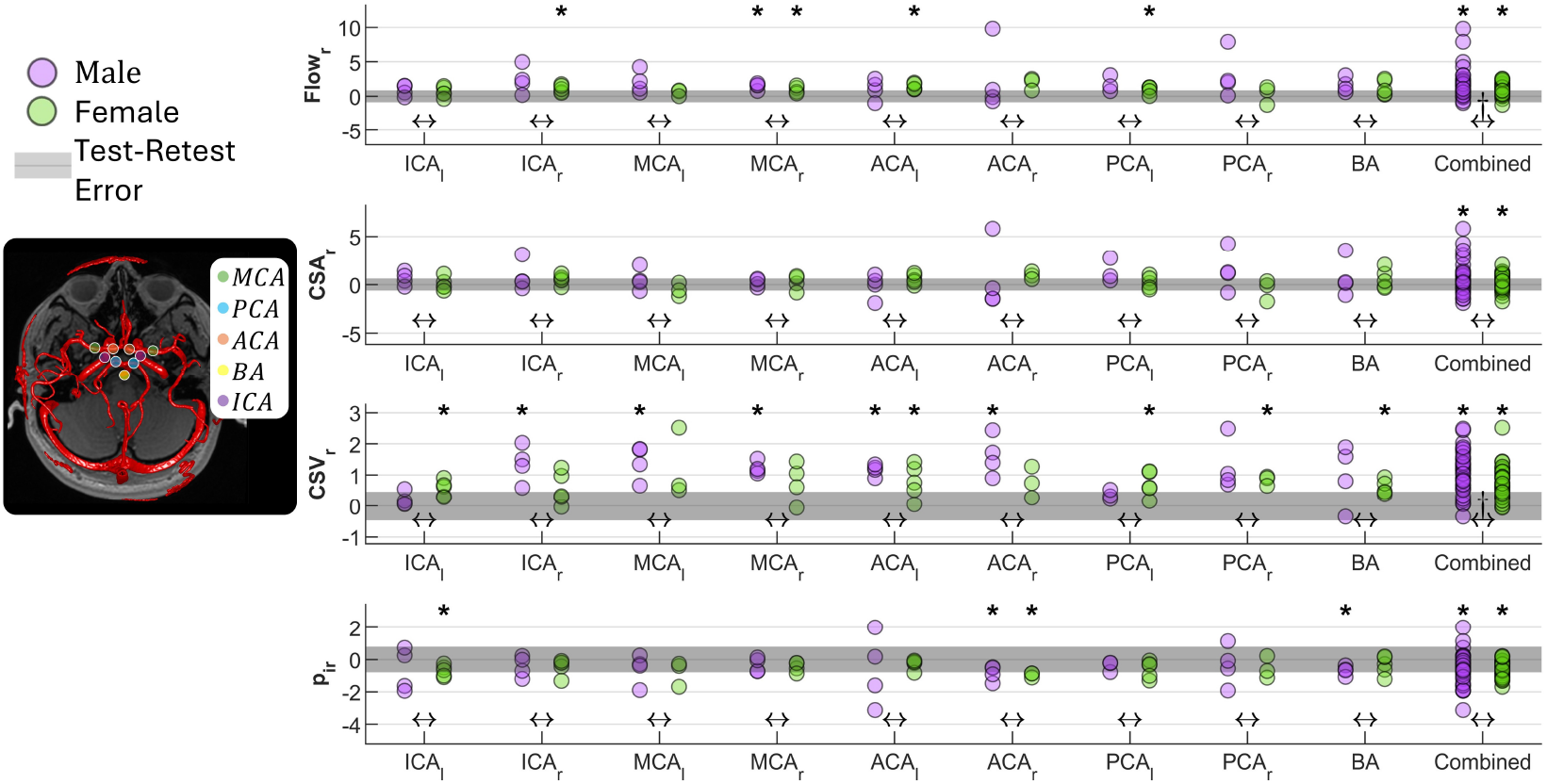
Vessel-specific cerebrovascular reactivities separated by sex. An asterisk above a group denotes a significant (p*<* 0.05) average reactivity, while a † between groups indicates a significant sex difference between reactivities. Symbols and abbreviations: internal carotid artery (ICA), middle cerebral artery (MCA), anterior cerebral artery (ACA), posterior cerebral artery (PCA), basilar artery (BA), cross-sectional area (CSA), cross-sectional velocity (CSV), pulsatility index (*p*_*i*_).

The most responsive vessel-specific index was CSV_*r*_, which had the largest number of confidently identifiable reactivities outside the test-retest limits, with a responsiveness of 76.7%. However, CSV_*r*_ also had the lowest dynamic range of ±0.6 (%Δ*/*%Δ), which was maximum for Flow_*r*_ with ±1.65 (%Δ*/*%Δ).

#### 3.3.2 Regional Reactivity

No consistent differences in ICA versus BA reactivities were identified for any index, and most regional CVRs were non-significant (see Figure 3). Both female ICAs and male RICA had significant CSV_*r*_, the male RICA also had a significant CSA_*r*_, and there were sex differences in the LICA CSA_*r*_. Regarding the combined data, the male CSA_*r*_ was significant and significantly higher than that for the female group, which also had an unreactive CSA_*r*_. Overall 21/27 regions dilated significantly. Both male and female groups showed a significant CSV_*r*_ with no sex differences observed. Overall, 25/27 regional velocities increased. None of PWV, *p*_*t*_, and *p*_*d*_ had significantly positive or negative CVR indices. PWV increased in 11*/*22 regions, *p*_*t*_ increased in 7*/*27 regions, and *p*_*d*_ increased in 12*/*27 regions. PWV and *p*_*t*_ had comparable CVRs and significantly larger standard deviations than any other index, *≈ ±*2.35 (%Δ*/*%Δ) versus the next largest of ±1.65 (%Δ*/*%Δ). PWV showed the highest responsiveness of regional indices of 86.4%.

**Figure 3.**
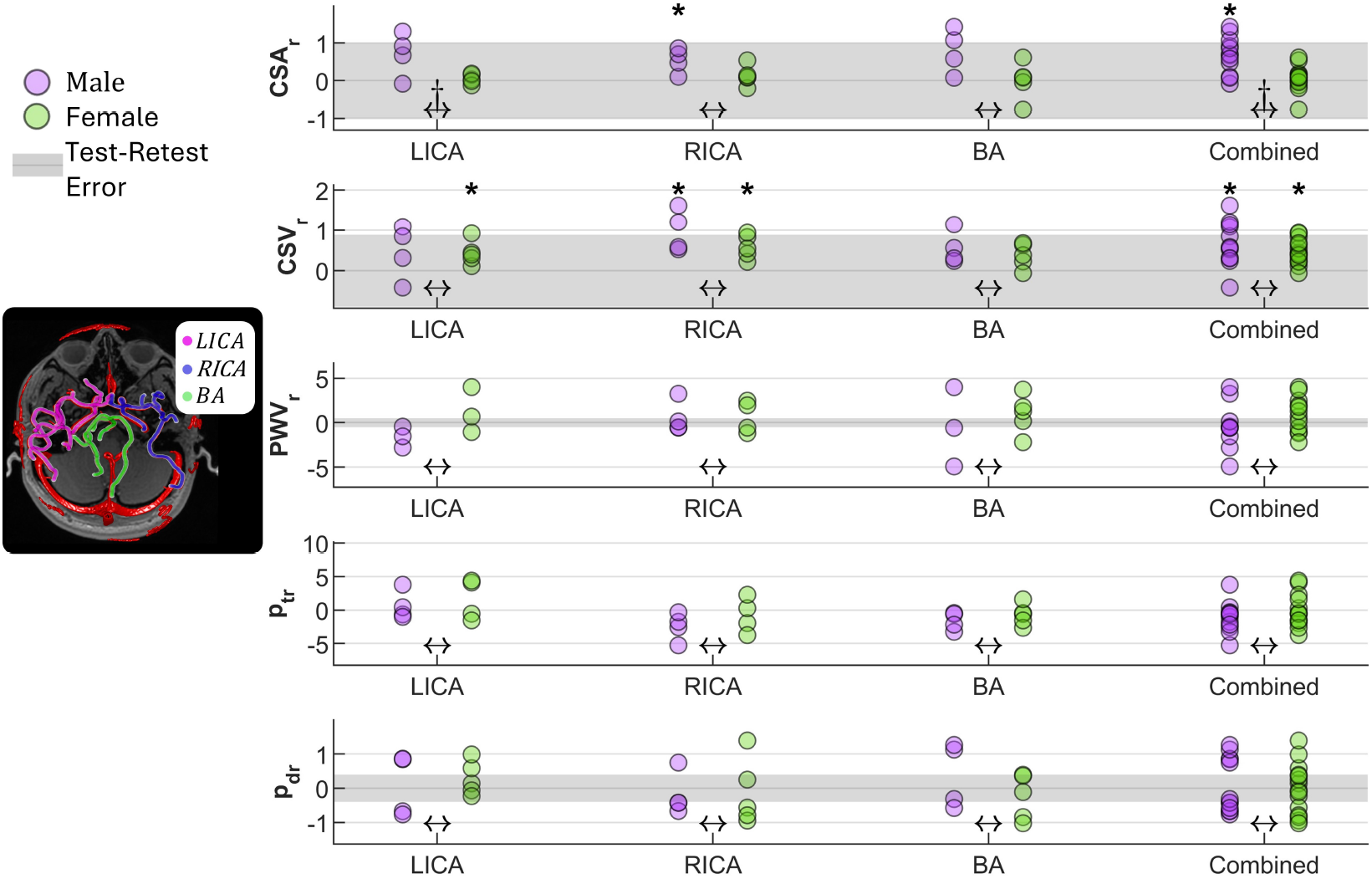
Regional cerebrovascular reactivities separated by sex. An asterisk above a group denotes a significant (p*<* 0.05) average reactivity, while a † between groups indicates a significant sex difference between reactivities. Symbols and abbreviations: left internal carotid artery (LICA), right internal carotid artery (RICA), basilar artery (BA), cross-sectional area (CSA), cross-sectional velocity (CSV), pulse wave velocity (PWV), pulsatility transmission (*p*_*t*_), pulsatility damping (*p*_*d*_).

#### 3.3.3 Global Reactivity

At global levels and including other cardiorespiratory measures, males exhibited significant positive reactivities in 7*/*10 indices (CBF, CSA, CSV, HR, P_*ET*_ CO_2_, SBP, DBP) with the exception of PWV, *p*_*t*_, and *p*_*d*_ (see Figure 4).

**Figure 4.**
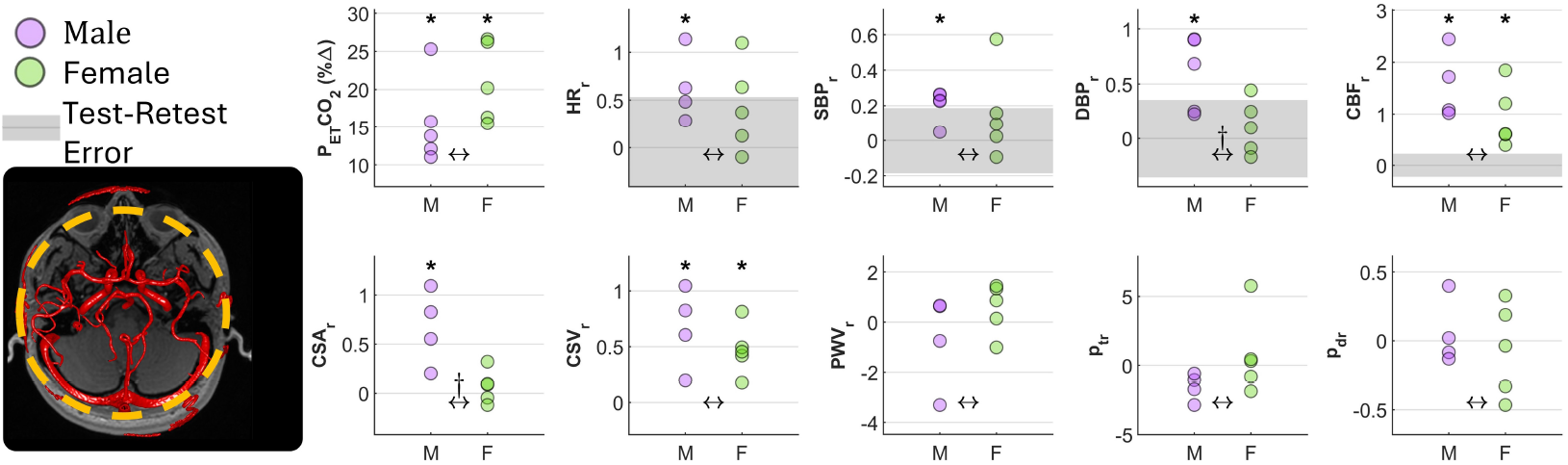
Global cardiorespiratory and cerebrovascular reactivities separated by sex. An asterisk above a group denotes a significant (p*<* 0.05) average reactivity, while a † between groups indicates a significant sex difference between reactivities. Symbols and abbreviations: partial pressure of end-tidal carbon dioxide (P_*ET*_ CO_2_), heart rate (HR), systolic blood pressure (SBP), diastolic blood pressure (DBP), cerebral blood flow (CBF), cross-sectional area (CSA), cross-sectional velocity (CSV), pulse wave velocity (PWV), pulsatility transmission (*p*_*t*_), pulsatility damping (*p*_*d*_).

Females exhibited only 3*/*10 significant reactivities (DBP, CSV, P_*ET*_ CO_2_). Although there was a large difference in the number of significantly reactive indices, only 3*/*10 indices had significant differences between the sexes; females had significantly lower CSA and SBP reactivities, and higher P_*ET*_ CO_2_ reactivity. Total CBF_*r*_ was the most responsive with a 100% responsiveness.

All combined reactivity data at all levels and confident responses are summarised in Table 2. The covariate analysis is reported in Supplementary Note 2.

**Table 2:** Combined index reactivity means and standard deviations assessed during hypercapnia and the number of confident responses outside test-retest error. An asterisk indicates a statistically significant (*p <* 0.05) response, and a † indicates a statistically significant sex difference. Symbols and abbreviations: male (M), female (F), cross sectional area (CSA), cross sectional velocity (CSV), pulsatility index (*p*_*i*_), pulse wave velocity (PWV), pulsatility transmission (*p*_*t*_), pulsatility damping (*p*_*d*_), partial pressure of end-tidal carbon dioxide (P_*ET*_ CO_2_), heart rate (HR), systolic blood pressure (SBP), diastolic blood pressure (DBP), cerebral blood flow (CBF)

| Vessel-Specific Reactivity |  |  |  |  |
| --- | --- | --- | --- | --- |
|  | Combined M | Combined F | Combined M&F | Confident Responses |
| Flow<sub>r</sub> | 1.85 ± 2.16* | 1.01 ± 0.79*<sup>†</sup> | 1.41 ± 1.65* | 63.0%(46/73) |
| CSA<sub>r</sub> | 0.67 ± 1.63* | 0.26 ± 0.76* | 0.46 ± 1.26* | 48.0%(35/73) |
| CSV<sub>r</sub> | 1.09 ± 0.67* | 0.72 ± 0.49*<sup>†</sup> | 0.89 ± 0.6* | 76.7%(56/73) |
| $p_{ir}$ | -0.55 ± 0.96* | -0.54 ± 0.47* | -0.54 ± 0.74 | 34.2%(25/73) |
| Regional Reactivity |  |  |  |  |
|  | Combined M | Combined F | Combined M&F | Confident Responses |
| CSA<sub>r</sub> | 0.68 ± 0.47* | 0.06 ± 0.31<sup>†</sup> | 0.33 ± 0.50* | 11.1%(3/27) |
| CSV<sub>r</sub> | 0.66 ± 0.55* | 0.46 ± 0.29* | 0.55 ± 0.43* | 22.2%(6/27) |
| PWV<sub>r</sub> | -0.40 ± 2.59 | 0.92 ± 1.96 | 0.32 ± 2.31 | 86.4%(19/22) |
| $p_{tr}$ | -1.15 ± 2.20 | -0.02 ± 2.49 | -0.56 ± 2.38 | - |
| $p_{dr}$ | 0.08 ± 0.80 | -0.03 ± 0.72 | 0.02 ± 0.74 | 70.4%(19/27) |
| Global Reactivity |  |  |  |  |
|  | M | F | M&F | Confident Responses |
| $P_{ET}CO_2$ (%$\Delta$) | 15.60 ± 5.71* | 20.97 ± 5.29*<sup>†</sup> | 18.29 ± 5.91* | 100%(9/9) |
| HR<sub>r</sub> | 0.63 ± 0.36* | 0.43 ± 0.47 | 0.52 ± 0.41* | 44.4%(4/9) |
| SBP<sub>r</sub> | 0.21 ± 0.09* | 0.15 ± 0.26 | 0.18 ± 0.18* | 55%(5/10) |
| DBP<sub>r</sub> | 0.59 ± 0.34* | 0.11 ± 0.25<sup>†</sup> | 0.35 ± 0.38* | 40%(4/10) |
| CBF<sub>r</sub> | 1.57 ± 0.66* | 0.93 ± 0.60* | 1.21 ± 0.68* | 100%(9/9) |
| CSA<sub>r</sub> | 0.67 ± 0.38* | 0.07 ± 0.17<sup>†</sup> | 0.33 ± 0.41* | - |
| CSV<sub>r</sub> | 0.67 ± 0.36* | 0.47 ± 0.23* | 0.56 ± 0.29* | - |
| PWV<sub>r</sub> | -0.69 ± 1.86 | 0.56 ± 1.01 | 0.00 ± 1.50 | - |
| $p_{tr}$ | -1.55 ± 0.99 | 0.77 ± 2.94 | -0.26 ± 2.49 | - |
| $p_{dr}$ | 0.05 ± 0.24 | -0.06 ± 0.34 | -0.01 ± 0.29 | - |

## 4 Discussion

The ability to modulate CVT, evaluated through stimulus-induced CVR, can lead to a better understanding of vascular regulation in cerebral pathologies. In this work, CVR indices were measured using 4D flow MRI at different levels, from vessel-specific to global assessments. As previously identified, flow and velocity increased in response to hypercapnia. Regional and global CSA analysis identified positive increases that were not clear at the vessel-specific level. Remarkably, the region-based reactivity of PWV, *p*_*t*_, — analysed for the first time here — increased in some cases and decreased in others. The PWV observation indicates that vasorelaxation in the main cerebral arteries may not always occur during hypercapnia as commonly assumed. Finally, at all levels and for almost all indices, there were consistent trends that CVR was lower in females compared to males.

### 4.1 Multi-Level CVR Integration With Previous Findings

The vessel-specific circle of Willis blood flow trends are generally consistent with previously reported data that used 4D flow MRI [30], [31], but each study, including the current, measured reactivity differently and used different CO_2_ concentrations, which confounds direct comparisons. Important consistencies are that flow reactivity was positive, and that females had lower reactivity than males. On the global flow level, we observed that normocapnic volume-normalised CBF was higher in young females, consistent with other reports [38], [39], and that young females have lower global CBF reactivity to hypercapnia [31].

To the best of our knowledge, only at the vessel-specific levels, have area and velocity reactivities have been explored [9]. In the present study, their regional and global averages have also been evaluated. This identified a consistently positive and greater male CSV reactivity at all studied levels, whereas such consistency was not observed for CSA. On the vessel-specific level, reactivity was minor, with many vessels constricting in males and females. At the regional level, females were non-reactive, while males were reactive. This may identify sex and spatially heterogeneous CVR, with dilation on average in men; however, with finer resolution, several studies have demonstrated that similarly sampled regions of the MCA and ICA dilate [40]–[42]. The discordance is likely to demonstrate an increase in the signal-to-noise ratio of the region-level evaluation. Vessel-specific analysis is prone to co-location error, and at these resolutions a single voxel can represent a 20% change in diameter [43], with each error incorrectly indicating contraction or dilation. By evaluating the hemodynamics of hundreds of locations to produce a representative index, regional metrics can reduce the impact of these segmentation and co-location errors. Having already demonstrated the ability to reveal physiological trends in pulsatility [5], an improvement in CSA signal-to-noise ratio is also likely. The CSA sex differences identified here have not been explored previously, so comparisons are not possible. However, these results are consistent with the notion that females have lower CVR, as reported previously [25]. The dilation identified in this work also reaffirms the notion that male velocity-based CVR is underestimated [31], [41], and further emphasises the need for caution when interpreting velocimetry studies, such as transcranial Doppler (TCD) studies that assumes vascular CSA remains constant [9].

The other global systemic reactivities reported here are consistent with reported values for HR [30], [31], [44] and blood pressure [19]–[21], [30], [31], including a greater increase in DBP in males [22]. The P_*ET*_ CO_2_ increased during hypercapnia, as expected [9], although other studies do not regularly report a larger female P_*ET*_ CO_2_ increase in response to hypercapnia as demonstrated in this work. This inconsistency is likely attributable to differences in the duration of hypercapnia. In other 4D flow reactivity studies [30], [31], P_*ET*_ CO_2_ data were not reported in a sex-specific manner. TCD studies have investigated sex differences in P_*ET*_ CO_2_, and reported no difference [45]. However, typical exposure in TCD studies is only a few minutes compared to the prolonged exposure in our study (*≈* 15 minutes) which possibly changed the gas dynamics, including allowing for CO_2_ normalisation. Lastly, a decreased *p*_*i*_ was observed during hypercapnia, in agreement with other literature [31], [46].

### 4.2 Interpreting Novel Pulsatility-Based and PWV Index Reactivity

As all standard processing was consistent with the literature, our data are most likely a good representation of a young normative cohort. Thus, the novel findings related to PWV, *p*_*t*_, and *p*_*d*_ discussed next are highly likely to also be representative, although the sample size is relatively small. This is a completely new approach to evaluate vascular health, where the following discussion will provide initial interpretation, but further work is needed for confirmation.

#### 4.2.1 Increased Dynamic Range

Neither PWV, *p*_*t*_, nor *p*_*d*_ showed a significant preference for positive or negative reactivity to hypercapnia, but they were responsive. In particular, PWV and *p*_*t*_ had the largest CVR standard deviations of 2.31 and 2.38 (%Δ*/*%Δ). This dynamic range is advantageous for an index to stratify pathology. Additionally, greater CVR variability may suggest that these regional indices provide stronger identification of physiological differences between participants. Although the target cohort was narrow in age and excluded co-morbidities, these participants have genetic and developmental differences from approximately 30 years of independent growth; therefore, differences are expected. This could mean that PWV and *p*_*t*_ are better suited to characterise subject-specific vascular function.

#### 4.2.2 Interpreting Positive and Negative Regional CVRs

In addition to the benefit of dynamic range, while all other indices generally increased in response to hypercapnia, PWV, *p*_*t*_, and *p*_*d*_ showed increases and decreases during hypercapnia beyond test-retest variability.

For *p*_*t*_ and *p*_*d*_, a decrease may reflect improved damping of the pulse wave. However, it was previously reported that *p*_*t*_ and *p*_*d*_ were negatively correlated with HR and CBF [5], both of which increased in response to hypercapnia. Therefore, we consider a *p*_*t*_ and *p*_*d*_ decrease as the non-reactive response (i.e., covariates can explain the difference). This was more common for males (p*<* 0.06) and, therefore, may not reflect a change in CVT. The cases for which *p*_*t*_ increased, particularly in females, identify a unique and unexpected change (the same as for *p*_*d*_). This could reflect a change in peripheral resistance via an increase in distal constriction and potentially a change in sympathetic activity. Interpreted as a risk factor for pulse wave encephalopathy, an increase in *p*_*t*_ is an unhealthy response that suggests increased pressure pulsatility in more sensitive tissues, consistent with increased female incidence of cerebrovascular disease [47]–[49]. A study on *p*_*d*_ hypercapnic reactivity of the extracranial ICA to intracranial MCA using Doppler ultrasound also reported a similar trend to overall improving damping (unreported sex differences or frequency of positive or negative reactivities) [50].

Turning attention to PWV, which has a positive correlation with HR and CSA [8], [37], both of which increased during hypercapnia, the non-reactive (covariate explained) response is for PWV to increase. This was more often observed for females, again implicating diminished CVR. A clear strength of measuring PWV is that it directly reflects CVT, where other indices can only infer a change in CVT. Thus, a decrease is generally interpretable as relaxation in CVT. However, when PWV increases, the cause is uncertain, it could be influenced by covariates (HR, CSA), low vascular chemosensitivity, or indicate competition with other mechanisms. The sympathetic response of females to hypercapnia was recently reported to be greater than that of males [51]. Therefore, increased PWV may be due to perivascular innervation and the CO_2_ sympathetic response outcompeting CVT relaxation in the imaged vasculature. This hypothesis also aligns with the minimal CSA reactivity we observed and suggests that downstream vascular dilation may be responsible for the increase in flow in females.

Regardless of the physiological cause for the increases and decreases in PWV, *p*_*t*_, and *p*_*d*_, this positive/negative variability provides additional physiological insight. Current hypercapnia-induced CVR indices (flow, CSV) are expected to be only positive. As one of the goals of measuring CVR is to assess vascular health and prognosis risk, an important step is to interpret the CVR values. When all values are of the same sign, a higher value may be beneficial in a prognostic sense, but even a minor reactivity still suggests some function. Prognostically, negative changes in PWV, *p*_*t*_, and *p*_*d*_ are always interpretable as haemodynamically healthy responses, while an increase is a clear indication of impaired CVR and the increased risk of pulse wave encephalopathy in situations where hypercapnia may occur, such as aligning with sleep apnoea and risk of dementia [52].

### 4.3 The Influence of Sex Hormones and Blood Composition

Both males and females demonstrated many dependencies on circulating sex hormones and blood composition in sex-dependent ways (Supplementary Note 2). The strongest trends identified in normocapnia are that testosterone was negatively correlated with PWV in males, and that testosterone and oestradiol influence P_*ET*_ CO_2_ in females. During hypercapnia, the SBP response was strongly regulated by testosterone. Mechanistic interpretation of the influence of sex hormones and blood-based factors is a matter for future CVR studies with larger samples, for which there appears to be value in their concurrent collection.

### 4.4 Limitations

Although it is clear that sample size is the largest limitation in this study, the consistency of the recorded responses with literature and trends that were repeated at several levels suggest that the novel reactivities of PWV, *p*_*t*_, and *p*_*d*_ were interpretable and representative. With regard to the reactivity index, we assessed relative reactivities to compare information between different indices. Absolute reactivity could also have been reported, but trends have been shown to be generally preserved between the relative and absolute reactivity calculations [53].

Positioning of the participants is a known factor that can influence reactivity [45]. Many studies have placed participants in the supine position, so the blood pressure results collected in the lateral decubitus position may not reflect the supine measurements of other studies. However, the observed trends in blood pressure are consistent with the previous studies. Recently, the influence of diurnal variability on resting CBF and PWV [37] has been reported. Although we considered including time of day as a covariate, the regular sleep cycles of the participants were not recorded, and without cycle normalisation to account for their biological clock, using time of day is not relevant. Interestingly, if the diurnal variability was separate from the testosterone cycle (which is also diurnal [54]), it is expected to decrease testosterone correlations. Thus, for the correlations that we observed, such as between testosterone and SBP reactivity (Supplementary Note 2), these values may be insensitive to diurnal variability.

Testosterone and oestradiol can be converted into one another via the aromatase enzyme [55]. This hormone conversion is hypothesised to be regulated locally in the brain [56], and it may be that the blood tests did not reflect cerebral concentrations, which may explain the lack of hormone trends with most cerebral measurements, although this could also be limited by the sample size. Concerning the influence of the menstrual cycle, we acknowledge that standard protocol aims to test females during the follicular phase of the menstrual cycle [9]. This was not part of the HER study protocol and it was a coincidence that all females were within the follicular phase (indicated by blood analysis charts). So while our results suggest diminished female reactivity, further studies for other phases of the menstrual cycle are warranted.

It is possible that bias was introduced for the normocapnia covariate analysis of *p*_*t*_, *p*_*d*_, and PWV by averaging territories, since magnitudes can vary significantly between the ICA and BA regions [5], [8]. However, there were no statistically significant differences in the reactivity magnitudes between the vascular regions, so the average is most likely acceptable for CVR interpretations, which was the focus of this work. Future investigations with more cases would benefit from global level analysis with conclusive measurements in all 3 vessels during normocapnia. Regions could also be evaluated separately with larger sample sizes. We note that the *p*_*t*_ magnitudes were significantly different on the Siemens Healthineers 3T Vida Fit scanner versus GE Healthcare 3T Signa Premiere [5], indicating a possible MRI vendor/sequence dependence. In terms of haemodynamic accuracy, the generally negative *p*_*t*_ measured in [5] is consistent with previous reports on *p*_*d*_ (using MRI and TCD) [4], [48], suggesting that harmonisation of the index for the 3T Vida Fit might be needed. Finally, in this study, the test-retest errors for the regional CSA and CSV were estimated from data from one participant, and as such should be considered preliminary. The significant reactivity and sex differences observed here at the regional and global level are motivation for a more comprehensive study of test-retest variability to examine regional CSA and CSV.

### 4.5 Study strengths

One of the greatest strengths of this study was the specific measurement of hormones and brachial blood pressure in addition to cerebral imaging, which were each recorded within just 1.5 hours for each participant. This is a substantial benefit compared to other studies, for which different measurement sessions were separated by days [31] as well as time of day [57], which can introduce confounding factors, such as menstrual (day separation) or testosterone (time of day) cycles, in addition to sleep quality or stress, among other factors. Notably, in most studies, sex hormones were not explicitly measured. Another benefit of this study is the regional analysis that revealed significant area dilations in males, when compared to the analysis of vessels separately, as has been done previously (see [9] and references therein). We reiterate that this has implications for techniques that evaluate cerebrovascular reactivity using velocities instead of flows, and our data suggests that young male reactivities may be underestimated in velocity-based TCD studies. Finally, our flow-based ensembled PWV measurements, using the Siemens Healthineers 3T Vida Fit scanner in this study, were not statistically different to those measured for this age range in a previous study using a GE Healthcare 3T Signa Premiere scanner [8], providing evidence of sequence/scanner invariance, which has obvious clinical benefits.

## 5 Conclusions

CVR is promising in assessing vascular function for applications in neurodegenerative diseases. In this field, the common practice is to evaluate single-vessels, which is prone to noise with larger test-retest errors and difficult colocation for comparison between participants. This work demonstrated the response of robust region-based indices that provide a new view on vascular function and CVR, particularly for recently developed *p*_*t*_ and PWV indices. Evaluation of these new indices identified a significant increase in the dynamic range of CVR that should improve cohort stratification. At vessel-specific, regional, and global levels, sex differences and diminished female CVR were identified, supporting the hypotheses of CVT dysregulation in females, which may explain sex-based pathological incidence and outcomes. This finding, coupled with the more interpretable region-based positive and negative CVRs, further demonstrates the value of advanced 4D flow indices to study vascular function. Finally, this study presented evidence of haemodynamic coupling with hormonal, blood pressure, and blood composition influences that are not often measured in single studies, and this strongly motivates and justifies their concurrent measurement.

## Data Availability

The raw scan data are available upon reasonable request to G.D.M.T. after appropriate institutional data sharing and ethics agreements have been met.

## Acknowledgments

We gratefully acknowledge the support from the University of Auckland Centre for Advanced MRI, and LabPLUS for facilitating MRI and blood collection, respectively, Jiantao Shen for supporting the vascular segmentation, Sunil Cherian (ADInstruments) for advice on the analyser and equipment, Neale Gover (Apex Medical Ltd.) and Kevin Rudolph (Hans Rudolph inc.) for advice on the gas delivery system and purchasing, and Sharlene Pacaigue for grant administration and management.

## Author Contributions

S.D., K.B., J.P.F., G.D.M.T. conceived and designed the research; S.D., H.W., D.Z., G.Q. performed the experiments; S.D. analysed the experimental data; S.D., J.P.F., G.D.M.T. interpreted the results; S.D. prepared the figures; S.D. drafted the manuscript; S.D., H.W., D.Z., G.Q., M.P.N., K.B., J.P.F., G.D.M.T. edited and revised the manuscript; S.D., H.W., D.Z., G.Q., M.P.N., K.B., J.P.F., G.D.M.T. approved the final version of the manuscript.

