## Supplementary for "Regional and Global Analysis of Cerebrovascular Reactivity Using 4D Flow MRI"

ACCESSIBLE AS A DOI AFTER REVIEW

**Sergio Dempsey***
Auckland Bioengineering Institute
University of Auckland


**Harvey Walsh**
Faculty of Medical and Health Sciences
University of Auckland


**Debbie Zhao**
Auckland Bioengineering Institute
University of Auckland


**Gina Quill**
Auckland Bioengineering Institute
University of Auckland


**Martyn P. Nash**
Auckland Bioengineering Institute &
Department of Engineering Science
and Biomedical Engineering
University of Auckland


**Kelly Burrowes**
Auckland Bioengineering Institute
University of Auckland


**James P. Fisher**[1]
Faculty of Medical and Health Sciences
University of Auckland


**Gonzalo D. Maso Talou**[1]
Auckland Bioengineering Institute
University of Auckland

[1]*These authors contributed equally*

May 16, 2025

### 1 Supplementary Note 1: Preliminary Repeatability of 4D Flow Indices

Several estimates of test-retest repeatability were not available in literature, however, data from [1] was available to provide a preliminary estimate for several indices studied in this work. One participant had 5 back-to-back 4D flow scans at different k-space accelerations with coverage centred on the circle of Willis. In [1] it was shown that the time-varying amplitudes diminished with increasing acceleration, here we assume that would not affect the time-averaged mean cross sectional velocity and area assessed in this work. Additionally, relative pulsatility values should be unchanged as the amplitude decrease was independent of vessel size, so the variability of pulsatile damping should not be influenced [1]. Therefore, we processed these five scans to report the repeatability of these indices (shown in Figure 1) and identified

*Corresponding Author: 701/Level 7, 70 Symonds Street, Grafton, Auckland, 1010, NZ.

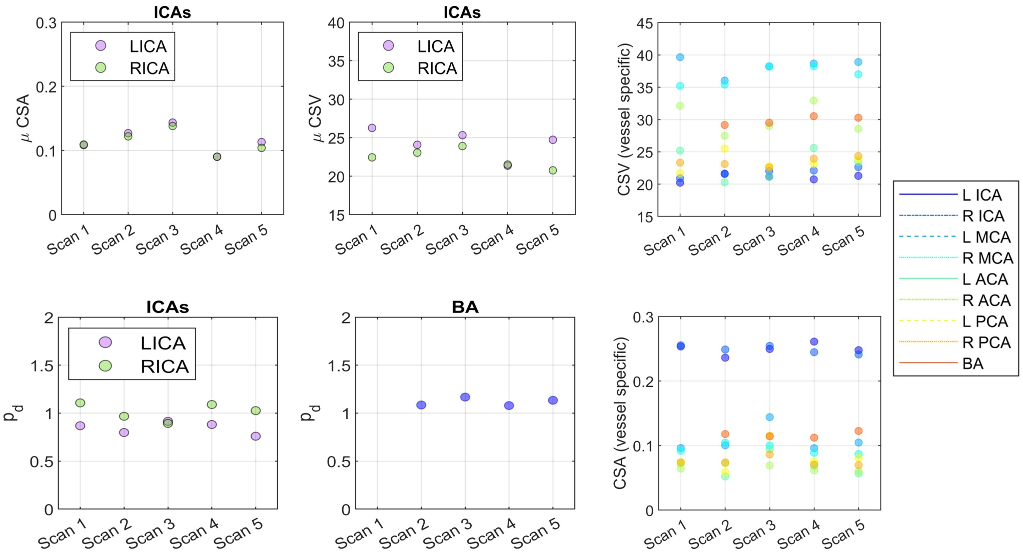


Figure 1: Repeatability of several indices after repeated 4D flow scans.

that these time averaged and relative values appeared to be repeatable. Unfortunately, the slice coverage was too low to confidently compute transmission or pulse wave velocity which has been shown to be sensitive to low slice coverage in [2].

### 2 Supplementary Note 2: Preliminary and Exploratory Global Covariate Analysis

As the scope of the covariates and indices considered in this work is much larger than conventional CVR covariates considered (that focus on one or two indices and typically only sex and age covariates), it is prudent to report the results in concise and interpretable ways, avoiding numerous plots with all possible combinations of data. As such, while correlations and linear models were fitted to compare all indices, a network graph plot was used to visualise only statistically significant relationships, shown in Figure 2. This graph identifies all significant correlations ($p < 0.05$). All raw data plots for significant correlations are presented in Figures 3 and 4.

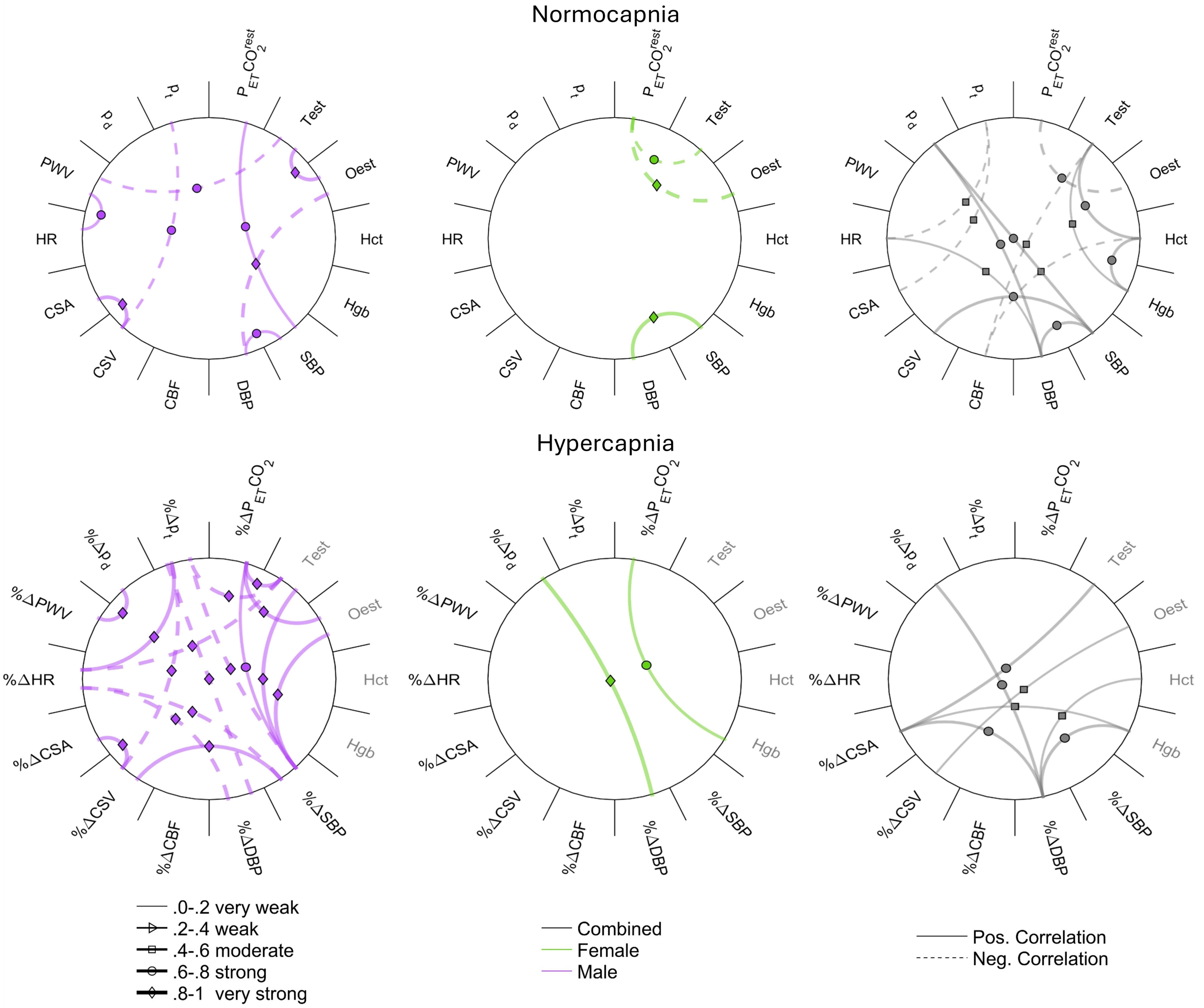


Figure 2: Network graph demonstrating significant correlations between reactivity indices. Abbreviations: cerebral blood flow (CBF), diastolic blood pressure (DBP), end-tidal carbon dioxide partial pressure ($P_{ET}CO_2$), hematocrit (Hct), haemoglobin (Hgb), heart rate (HR), mean cross-sectional area (CSA), mean cross-sectional velocity (CSV), oestradiol (oest), pulsatility transmission ($p_t$), pulse wave velocity (PWV), systolic blood pressure (SBP), testosterone (Test).

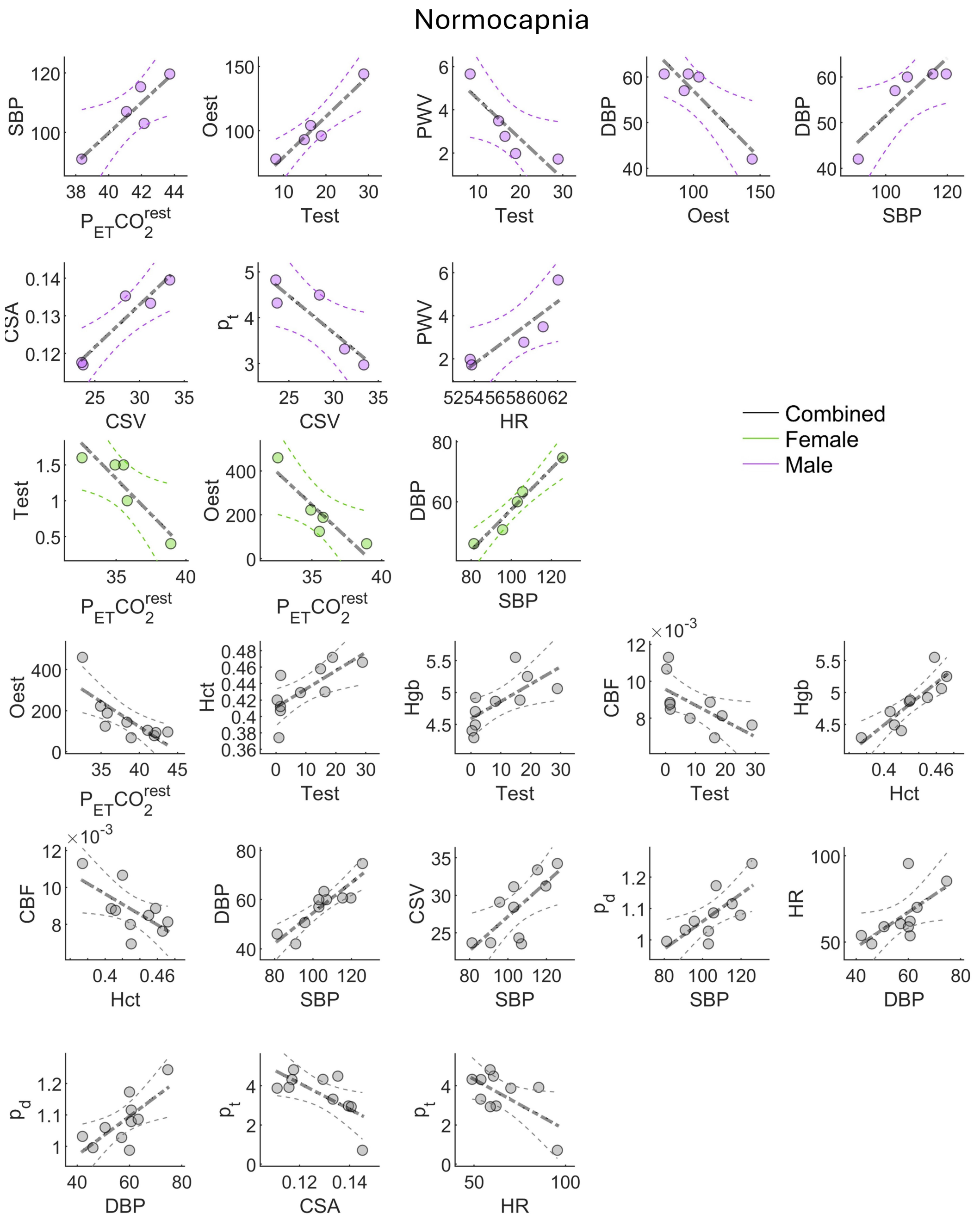


Figure 3: Raw data of significantly ($p < 0.05$) correlated indices for resting normocapnia. Abbreviations: Abbreviations: cerebral blood flow (CBF), diastolic blood pressure (DBP), end-tidal carbon dioxide partial pressure ($P_{ET}CO_2$), haematocrit (Hct), haemoglobin (Hgb), heart rate (HR), mean cross-sectional area (CSA), mean cross-sectional velocity (Vel), oestradiol (oest), pulsatility transmission ($p_t$), pulse wave velocity (PWV), systolic blood pressure (SBP), testosterone (Test).

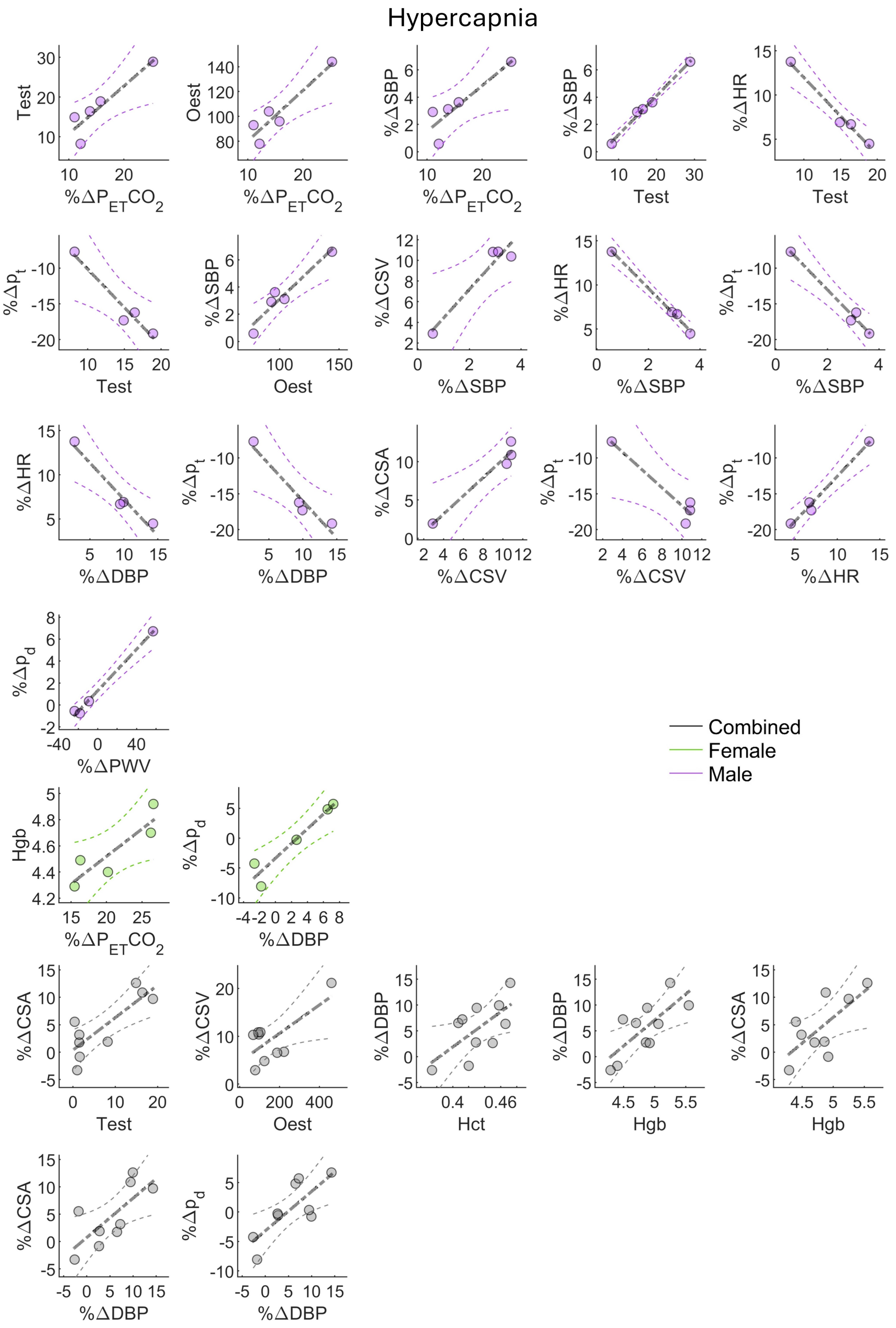


Figure 4: Raw data of significant ($p < 0.05$) correlated indices for hypercapnic reactivity. Abbreviations: Abbreviations: cerebral blood flow (CBF), diastolic blood pressure (DBP), end-tidal carbon dioxide partial pressure ($P_{ET}CO_2$), haematocrit (Hct), haemoglobin (Hgb), heart rate (HR), mean cross-sectional area (CSA), mean cross-sectional velocity (Vel), oestradiol (oest), pulsatility transmission ($p_t$), pulse wave velocity (PWV), systolic blood pressure (SBP), testosterone (Test).
